# Long-term effects of wildfire smoke exposure during early-life on the nasal epigenome in rhesus macaques

**DOI:** 10.1101/2021.09.10.459648

**Authors:** Anthony P. Brown, Lucy Cai, Benjamin I. Laufer, Lisa A. Miller, Janine M. LaSalle, Hong Ji

## Abstract

**Background:** Wildfire smoke is responsible for around 20% of all particulate emissions in the U.S. and affects millions of people worldwide. Children are especially vulnerable, as ambient air pollution exposure during early childhood is associated with reduced lung function. Most studies, however, have focused on the short-term impacts of wildfire smoke exposures. We aimed to identify long-term baseline epigenetic changes associated with early-life exposure to wildfire smoke. We collected nasal epithelium samples for whole genome bisulfite sequencing (WGBS) from two groups of adult female rhesus macaques: one group born just before the 2008 California wildfire season and exposed to wildfire smoke during early-life (n = 8), and the other group born in 2009 with no wildfire smoke exposure during early-life (n = 14). RNA-sequencing was also performed on a subset of these samples.

**Results:** We identified 3370 <u>d</u>ifferentially <u>m</u>ethylated <u>r</u>egions (DMRs) (difference in methylation ≥ 5% empirical p < 0.05) and 1 differentially expressed gene (*FLOT2*) (FDR< 0.05, fold of change ≥ 1.2). The DMRs were annotated to genes significantly enriched for synaptogenesis signaling, protein kinase A signaling, and a variety of immune processes, and some DMRs significantly correlated with gene expression differences. DMRs were also significantly enriched within regions of bivalent chromatin (top odds ratio = 1.46, q-value < 3 x 10^-6^) that often silence key developmental genes while keeping them poised for activation in pluripotent cells.

**Conclusions:** These data suggest that early-life exposure to wildfire smoke leads to long-term changes in the methylome over genes impacting the nervous and immune systems, but follow-up studies will be required to test whether these changes influence transcription following an immune/respiratory challenge.

## BACKGROUND

According to the National Interagency Fire Center, there were 50,477 wildfires (4.7 million acres burned) in the United States in 2019. In total, 212 million Americans lived in counties affected by wildfires in 2011 (1). These wildfires have contributed to levels of air pollution in the United States that have been linked to premature death (2–5). About 20% of all fine particulate emissions in the U.S. are from wildfire smoke, while half of all particulate matter less than 2.5 µm in diameter (PM_2.5_) in California resulted from wildfires (2). PM_2.5_ are especially harmful, as these particles are able to penetrate the respiratory system and the lungs (2). Exposure to these particles has been associated with asthma, bronchitis, lung cancer, and cardiovascular disease (3–5). Young children are especially vulnerable to these negative health effects, as studies have linked air pollution exposure in children to reduced lung function (6, 7), reduced height-for-age (8), increased blood pressure (9), and an increased risk of developing asthma and eczema (10). Most of these studies, however, focused on the short-term effects of exposures to wildfire smoke or polluted air and none have performed an unbiased assessment of gene pathways impacted by wildfire smoke exposure.

A cohort of rhesus macaques (*Macaca mulatta*) that were exposed in their first three months of life to a harsh wildfire season in 2008 in California was previously studied to understand some of the long-term effects of wildfire smoke exposure (11). Peripheral blood mononuclear cells (PBMCs) were cultured and challenged with either LPS or flagellin, and secretions of IL-8 and IL-6 were compared to macaques that were born in 2009 (PM_2.5_ and ozone levels were much lower in 2009 compared to 2008) (11). Lung function was also compared between exposed and control macaques. Compared to control macaques, wildfire smoke-exposed macaques had significantly reduced lung volume. Female wildfire-exposed macaques showed reduced production of IL-8 compared to controls, while male wildfire-exposed macaques showed reduced production of IL-6 compared to controls (11). This study implied that early-life exposure led to a difference in IL-8 and IL-6 production following an immune challenge, but it was still unclear to what degree these macaques exhibited baseline differences at the level of epigenetics and gene expression.

The epigenetic mark of DNA methylation has the potential to reflect past exposures with long-lived marks on genes, while the transcriptome reflects current levels of gene expression in a sampled tissue. To test the hypothesis that early-life wildfire smoke exposure would result in detectable epigenetic differences to gene pathways reflecting cellular function, we performed the integrated unbiased approaches of whole genome bisulfite sequencing (methylome) and RNA-sequencing (transcriptome) from nasal epithelial samples collected the same cohorts of female macaques examined a decade earlier for lung functions and immune responses. We identified a large number of genes associated with early-life exposure-related differential methylation involved in neuronal and immune signaling. In contrast, only one differentially expressed gene (*FLOT2*) was stably associated with early-life wildfire smoke exposure.

## RESULTS

### Exposure to wildfire smoke during infancy is associated with long-lasting changes to DNA methylation patterns in nasal epithelial cells

To test the effects of early-life wildfire smoke-exposure on methylation status throughout the genome, we performed whole genome bisulfite sequencing on nasal epithelial samples collected from 22 adult rhesus macaques in 2019 (8 born in 2008 and exposed to high levels of PM_2.5_ and ozone due to wildfires, 14 born in 2009 and therefore has relatively low levels of exposure to PM_2.5_ and ozone; Figure 1, Table 1). Though there were several shared exposures to high levels of wildfire smoke PM_2.5_ (> 35ug/m^3^, the 24-hour PM_2.5_ National Ambient Air Quality Standard) and ozone after the 2009 cohort was born (especially in the year of 2019), there was one high exposure event that only the 2008 cohort was exposed to in early-life (10 days above 35ug/m^3^, Figure 1, Table 1). We assessed 26,609,677 CpG sites and identified 3370 differentially methylated regions (DMRs) between exposed and non-exposed samples (Figure 2, empirical p < 0.05, differences in methylation>5%). The majority of these DMRs were hypermethylated in exposed animals (2899, ∼86%). A total of 114 (3.38%) of these DMRs were primarily located in CpG islands (12), 287 (8.52%) were located in CpG shores (0-2kb from island), 205 (6.08%) were located in CpG shelves (2-4kb from island), and 2764 (82.02%) were in the open sea (>4kb from island). This distribution was significantly different than expected by chance, with an enrichment towards CpG islands, shores, and shelves compared to regions assayed (Supplementary Figure 1). These 3370 DMRs were annotated to 2139 genes (Supplementary Table 1), of which 1852 genes were associated with DMRs hypermethylated in the exposed group, while 376 genes were associated with DMRs hypomethylated in the exposed group, and 89 genes were associated with both hypermethylated and hypomethylated DMRs (examples of DMRs shown in Figure 3). The DMRs were significantly more associated with promoters and exons than expected by chance, while they were less associated with intergenic regions than expected by chance (Supplementary Figure 2). The genes associated with DMRs as a whole were significantly enriched (FDR < 0.05) for 186 IPA canonical pathways, including *axonal guidance signaling*, *synaptogenesis signaling pathway*, *protein kinase A signaling*, *IL-15 production*, *CXCR4 signaling*, and *Th1 and Th2 activation pathway* (Figure 4, Supplementary Table 2).

**Figure 1:**
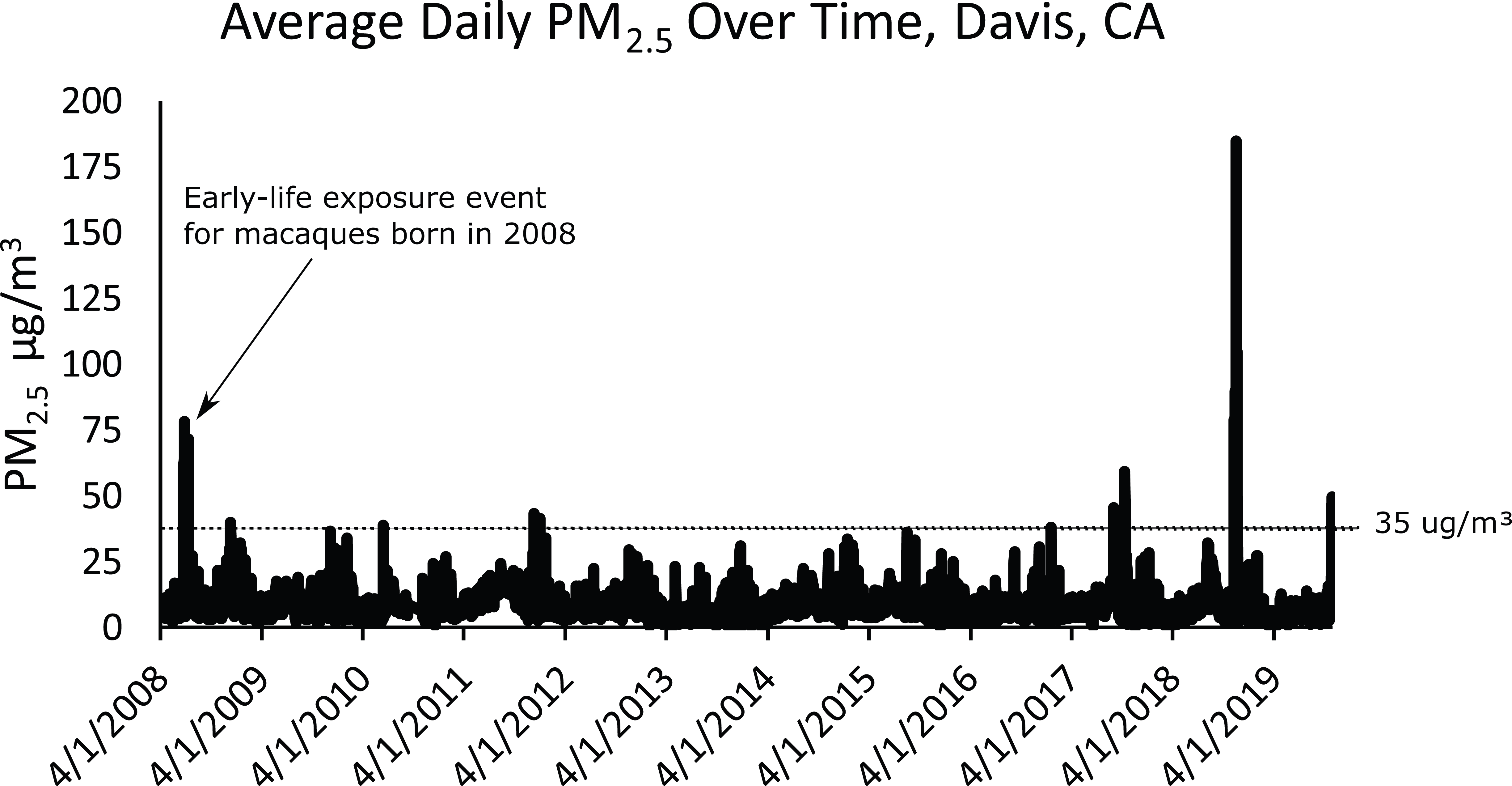
Average daily PM_2.5_ from April 2008 through October 2019 at the California Air Resources Board air monitoring station (site no. 57,577) located 2.7 miles southeast of the California National Primate Research Center on the University of California Davis campus. The dotted line at 35ug/m^3^ represent the 24-hour PM_2.5_ National Ambient Air Quality Standard. Note the arrow pointing to the early-life exposure event in macaques born in 2008. All other exposure events were shared between the two groups.

**Figure 2:**
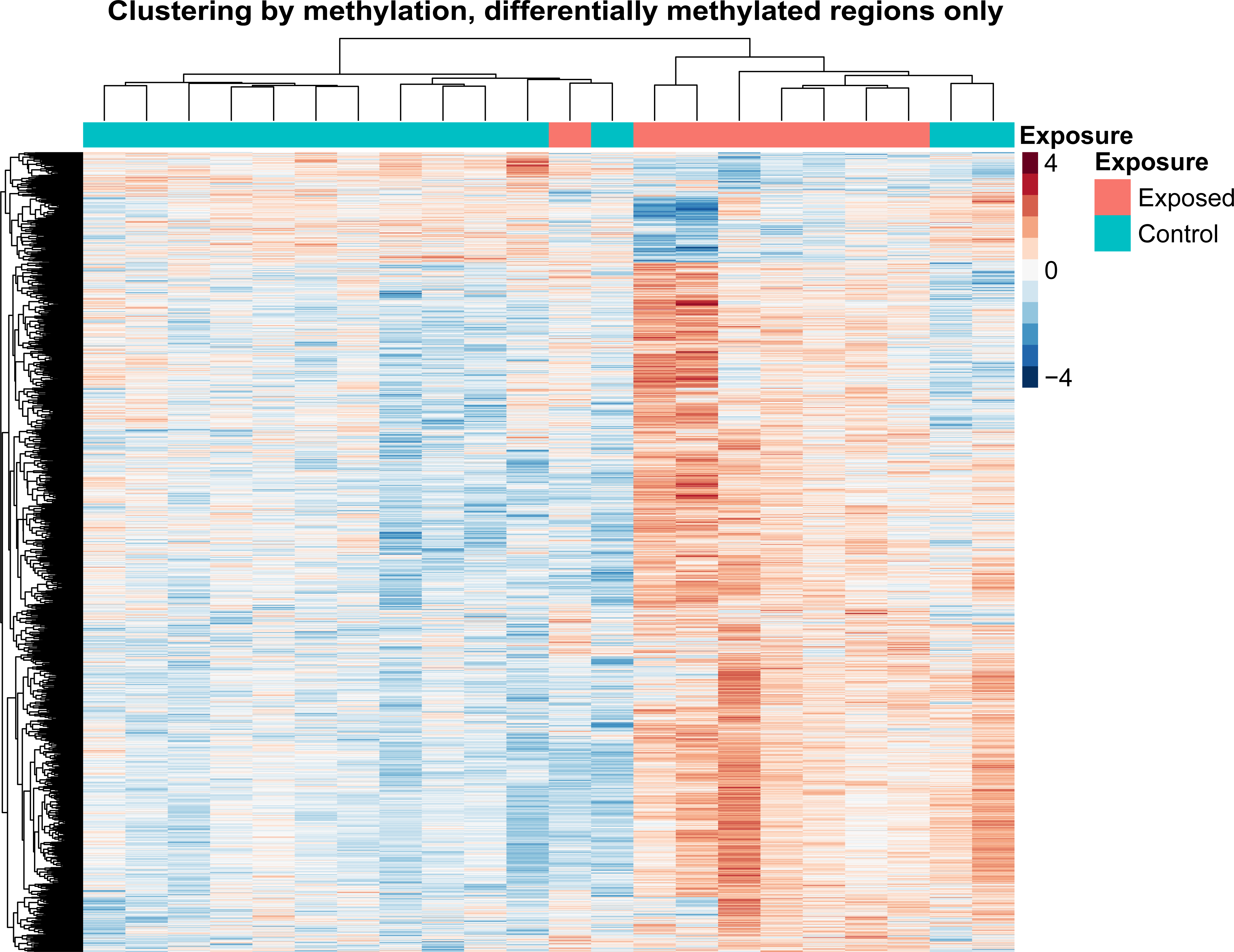
Heatmap showing sample clustering based on methylation. The heatmap includes only differentially methylated regions (DMRs). The heatmap was normalized on a per row basis for visualization, therefore the values on the scales are relative rather than absolute.

**Figure 3:**
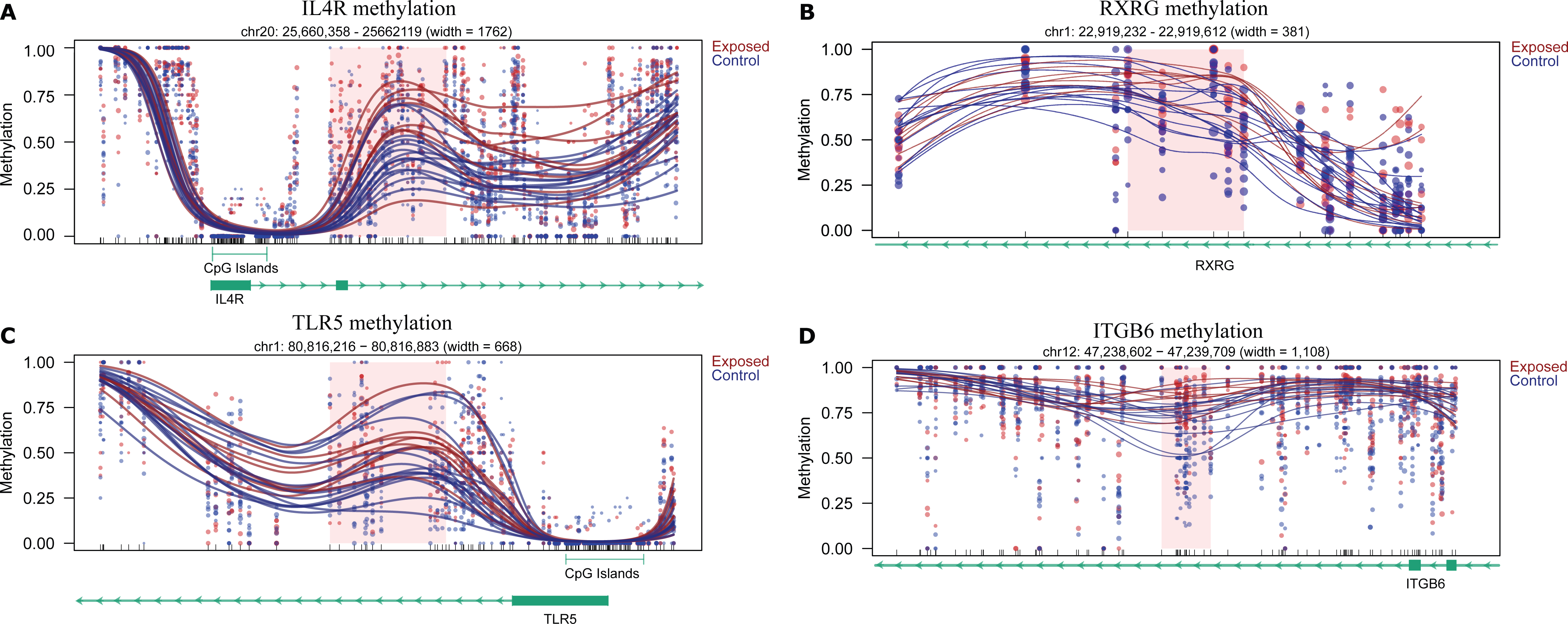
Examples of differentially methylated regions (DMRs) between rhesus macaques exposed in the first three months of life to wildfire smoke and those that were not. A) IL4R. B) RXRG. C) TLR5. D) ITGB6. Each dot represents the methylation percentage of one individual at one CpG site, while each line represents the smoothed average methylation level moving across the region. The red shaded boxes denote the specific DMR locations. Tracks for CpG islands (if present) or genes are included underneath each plot. For the gene tracks, a solid box indicates an exon, while the arrows indicate the direction of transcription.

**Figure 4:**
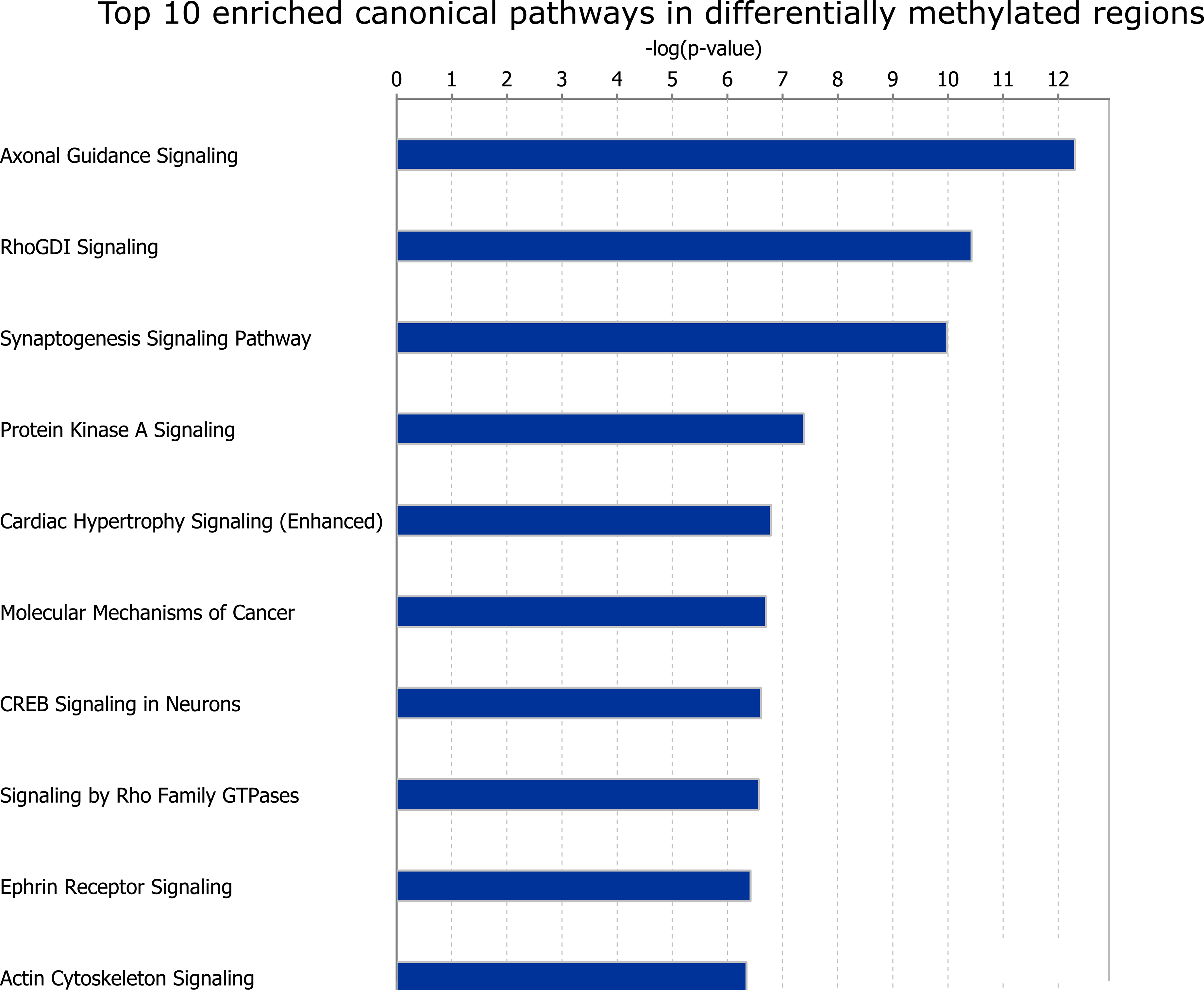
Enriched pathway analyses for differentially methylated regions (DMRs). Only the top ten (out of 186) enriched Ingenuity Pathway Analysis (IPA) canonical pathways are shown.

**Table 1:**
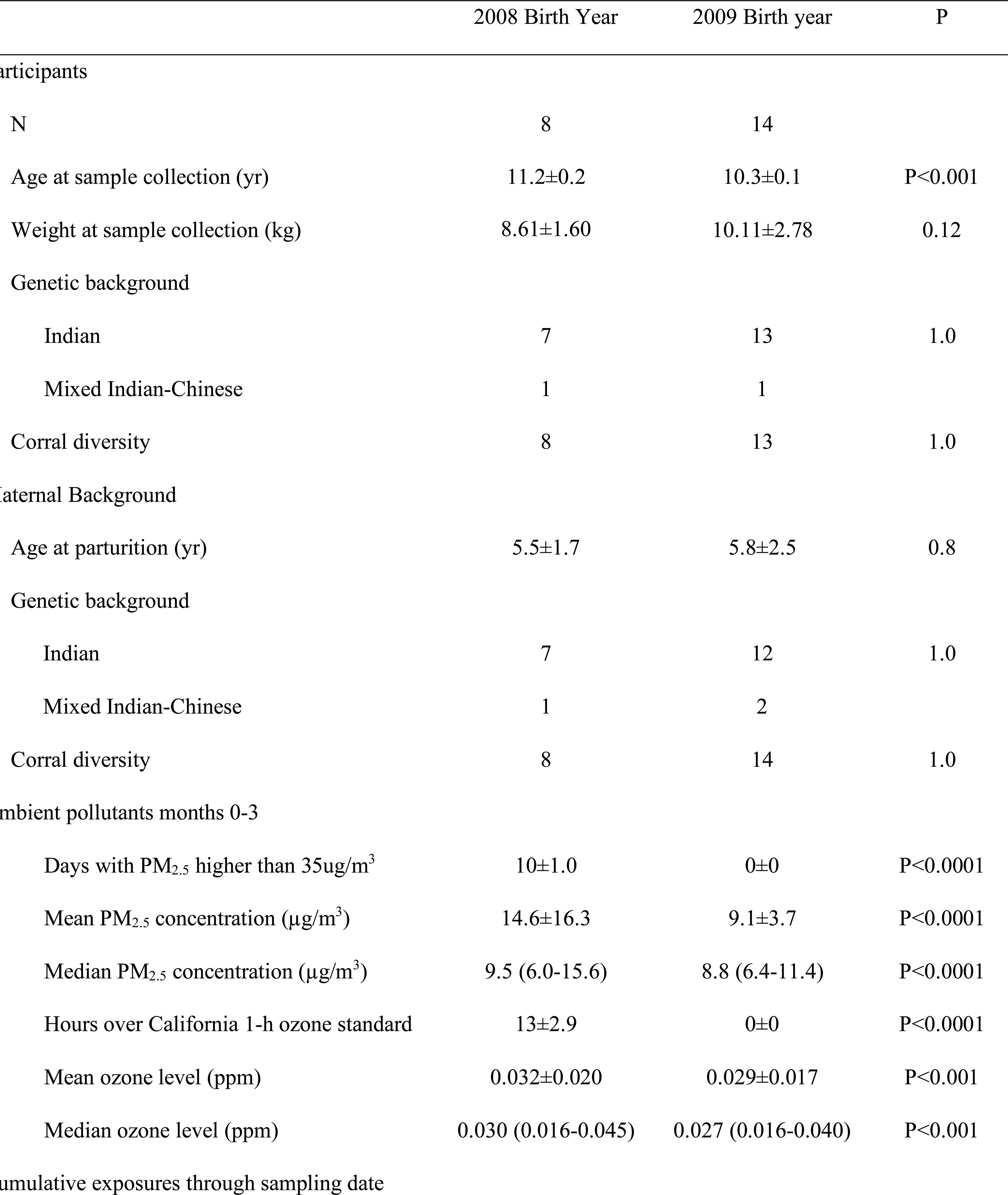
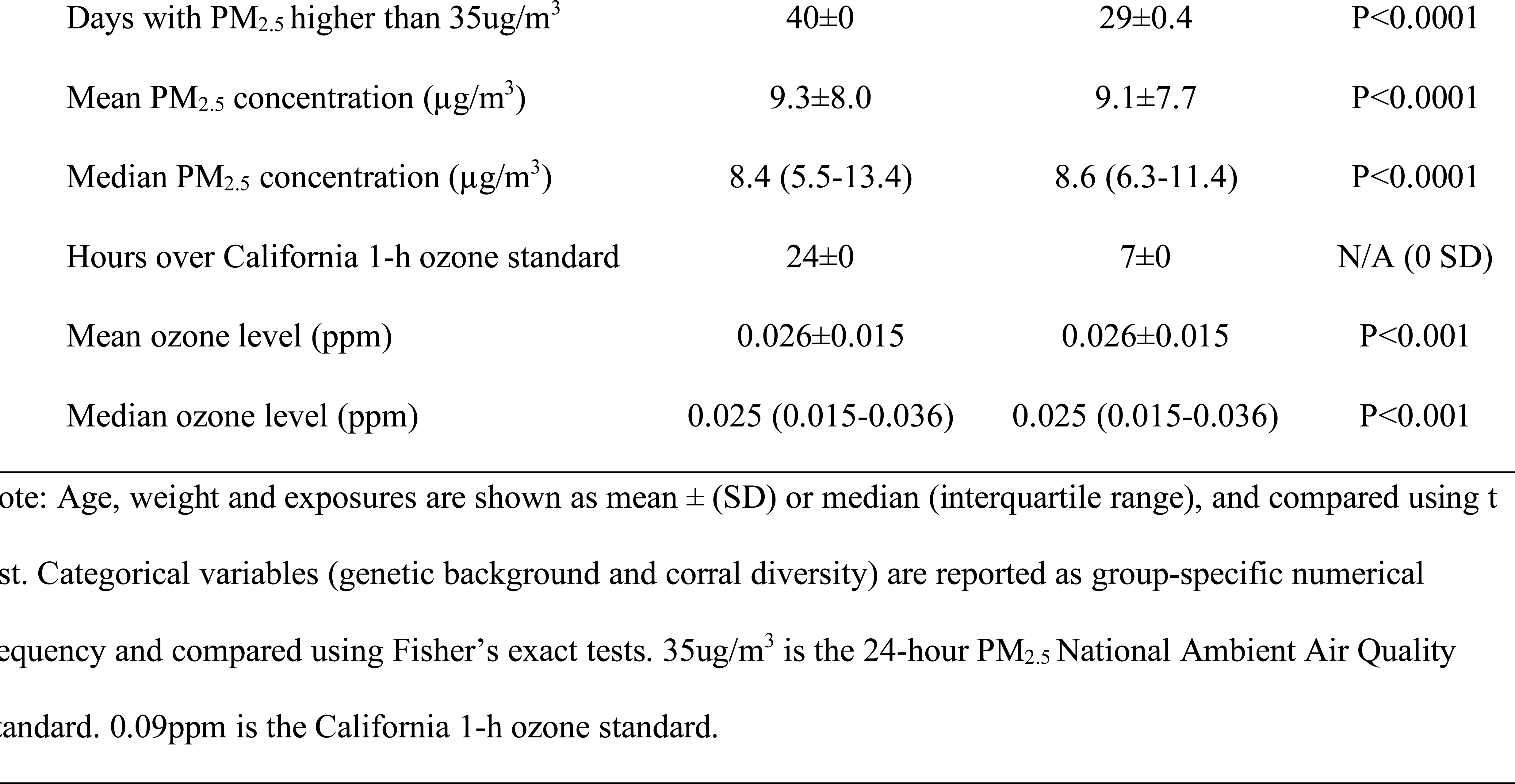
Demographic characteristics of animal populations

Genes associated with hypermethylated DMRs were enriched for 187 IPA pathways, 168 of which were also enriched in genes associated with DMRs as a whole. The 19 unique IPA pathways enriched in hypermethylated DMRs include *14-3-3-mediated signaling, LPS-stimulated MAPK Signaling,* and *NF-κB activation by viruses* (Supplementary Table 3). Genes associated with hypomethylated DMRs were enriched for 41 IPA pathways, 23 of which were also enriched in genes associated with DMRs as a whole. The 18 unique IPA pathways enriched in hypomethylated DMRs include *dermatan sulfate biosynthesis, xenobiotic metabolism PXR signaling pathway*, and *HOTAIR regulatory pathway* (Supplementary Table 4).

### Impact of wildfire smoke-associated DNA methylation changes on TF binding

As the binding of transcription factors (TFs) are often influenced by DNA methylation, we performed a HOMER analysis to determine whether any transcription factor binding sites were enriched in these wildfire smoke-associated DMRs (13). A total of 131 transcription factor motifs were enriched in all DMRs (q < 0.05; Supplementary Table 5). Eight of the top ten most highly enriched TF motifs are part of the bZIP TF family (shown in Table 2). When testing for TF binding site enrichment in only DMRs that were hypermethylated in exposed macaques, six of the top ten were part of the bZIP TF family, while none of the top ten enriched TF binding sites in hypomethylated DMRs were part of the bZIP TF family (five out of ten contained homeobox motifs). Interestingly, the TFs whose binding sites were most enriched in all wildfire smoke-associated DMRs were primarily unmethylated (Table 2) in other ChIP-seq datasets (14), so the differential methylation could theoretically have a large impact on transcription factor binding and expression (15). In support of this, DNA methylation generally inhibits binding of bZIP TF members to DNA (15, 16).

**Table 2:**

| Motif | unmethylated | partially methylated | methylated |
| --- | --- | --- | --- |
| Fra1/FOSL1 | 0.68 | 0.32 | 0.00 |
| Atf3 | 0.82 | 0.18 | 0.01 |
| JunB | 0.88 | 0.12 | 0.00 |
| BATF | 0.69 | 0.28 | 0.03 |
| Fra2/FOSL2 | 0.90 | 0.10 | 0.00 |
| AP-1/Jun | 0.81 | 0.19 | 0.00 |
| p63/TP53 | 0.46 | 0.35 | 0.19 |
| NF1-halfsite(CTF)/NFIA | 0.64 | 0.31 | 0.05 |

### Regions with hypomethylated DMRs are enriched for bivalent chromatin marks across tissue types

In order to understand the gene regulatory role of regions with wildfire smoke-DMRs, we searched for the enrichment of 15 pre-defined chromatin states across 127 epigenomes from multiple tissues and cell types in the Roadmap Epigenomics project (17). After converting the *M. maculatta* coordinates into human (hg38) coordinates and using LOLA, the DMRs as a whole were enriched for bivalent chromatin marks (top odds ratio for any mark = 1.46, q-value < 3 x 10^-6^; Figure 5A). Bivalent chromatin marks represent co-existing activating and repressing marks, which often silence key developmental genes while keeping them poised for activation in pluripotent cells (18). Hypomethylated DMRs seemed to drive this enrichment (top odds ratio for any mark = 2.05, q-value < 0.02; Figure 5B), though hypermethylated DMRs showed enrichment (top odds ratio for any mark = 1.51, q-value < 1 x 10^-6^) for bivalent chromHMM chromatin states as well (Figure 5C).

**Figure 5:**
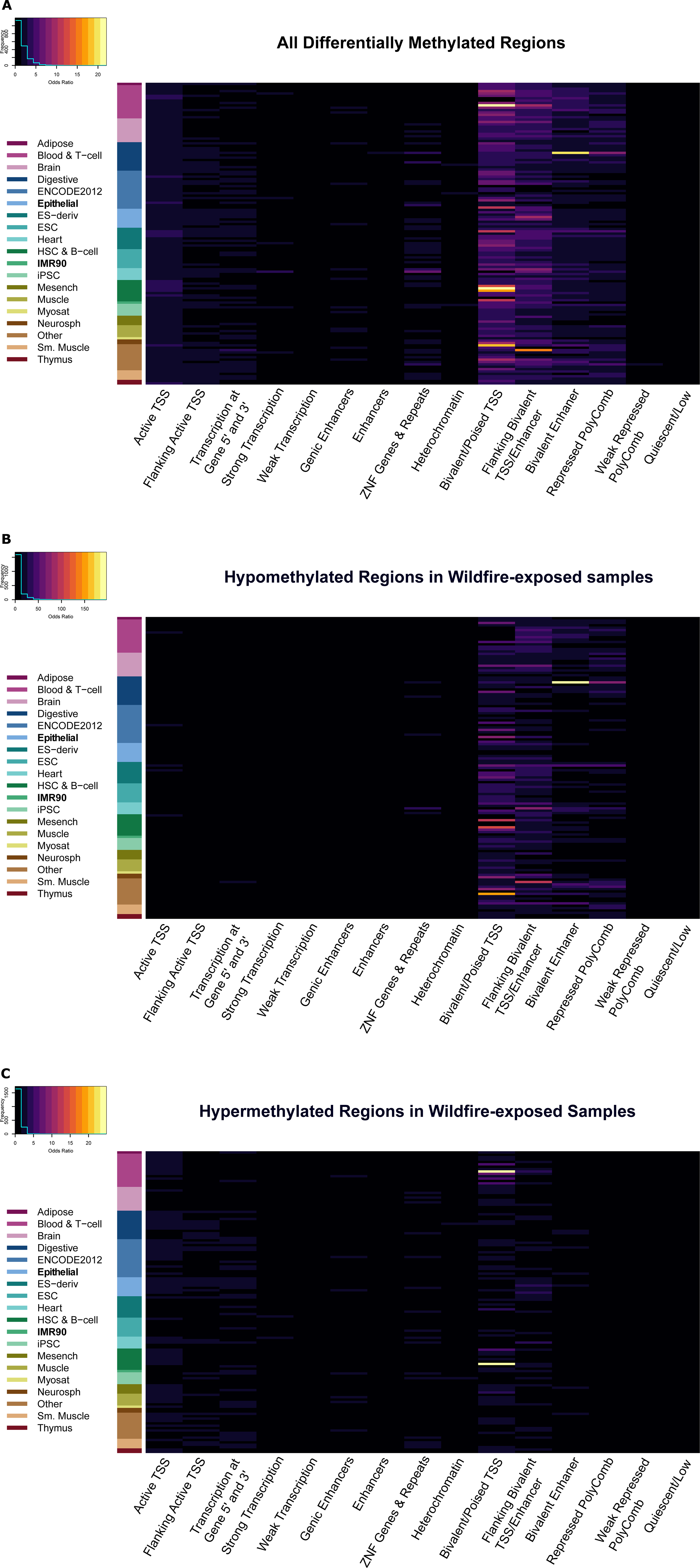
Enrichment in chromHMM (79) states in A) all differentially methylated regions (DMRs), B) DMRs that were hypomethylated in wildfire smoke-exposed macaques, and C) DMRs that were hypermethylated in wildfire smoke-exposed macaques. The rows in the plot represent different datasets from different cell types from the NIH Roadmap Epigenomics Consortium (88). Epithelial and IMR90 are highlighted in the plots, as these are the closest to the nasal epithelial samples in our current study.

### Early-life wildfire smoke exposure had a minimal effect on baseline genes expression levels

To determine whether early-life exposure to wildfire smoke leads to detectable differences in gene expression later in life, we performed RNA-sequencing on 15 female rhesus macaques (6 born in 2008 and exposed to wildfire smoke, 9 born in 2009 and not directly exposed to the 2009 California wildfires). A principal component analysis (PCA) and hierarchical clustering of all detected transcripts were performed to visualize how samples clustered based on expression (Supplementary Figure 3D). The top two principal components in a principal component analysis (PCA) explained 62% of the variation in the dataset. Exposed and non-exposed samples did not cluster separately in either the PCA or the hierarchical clustering analysis, implying no widespread transcriptomic difference between exposed and non-exposed individuals. After multiple hypothesis correction (FDR < 0.05, fold change ≥ 1.2; Supplementary Table 6), there was only one differentially expressed gene (*FLOT2*; Supplementary Table 6). None of the genes annotated to DMRs were significantly differentially expressed.

To identify co-expressed genes whose expression correlated with wildfire smoke-exposure status, we performed a weighted gene co-expression network analysis (WGCNA) (19). We identified 16 co-expressed modules using WGCNA. None of the modules were significantly associated with early-life exposure status (p < 0.05). The module that best correlated with exposure status was the purple module (p = 0.1; consisting of 585 genes, including IFI44, IFNA21, and IL24; Supplementary Figure 4, Supplementary Table 7). No genes in this module were significantly differentially expressed at an individual level, 19 genes were associated with significant DMRs, and two genes had significantly correlated methylation and expression. The genes in this module were enriched (FDR < 0.05) for 21 IPA pathways, including *EIF2 signaling*, *mTOR signaling*, *Th17 activation pathway*, and *interferon signaling* (Supplementary Table 8).

### Correlation of DNA methylation and gene expression differences resulting from wildfire smoke exposure during infancy

Out of the 2139 genes associated with DMRs, 2128 had enough corresponding expression data to evaluate the correlation between expression and methylation. To identify genes where differential methylation may be ultimately leading to differential expression, we calculated the spearman rank correlation between methylation and expression levels for genes that were associated with DMRs. In total, 172 genes were significantly correlated (spearman p-value < 0.05), with 76 genes showing a negative correlation and 96 showing a positive correlation between methylation and expression (Supplementary Table 9, two examples are shown in Figure 6). These 172 genes were enriched for 32 IPA pathway terms, including *leukocyte extravasation signaling, CCR5 signaling in macrophages,* and *MIF regulation of innate immunity* (Supplementary Table 10).

**Figure 6.**
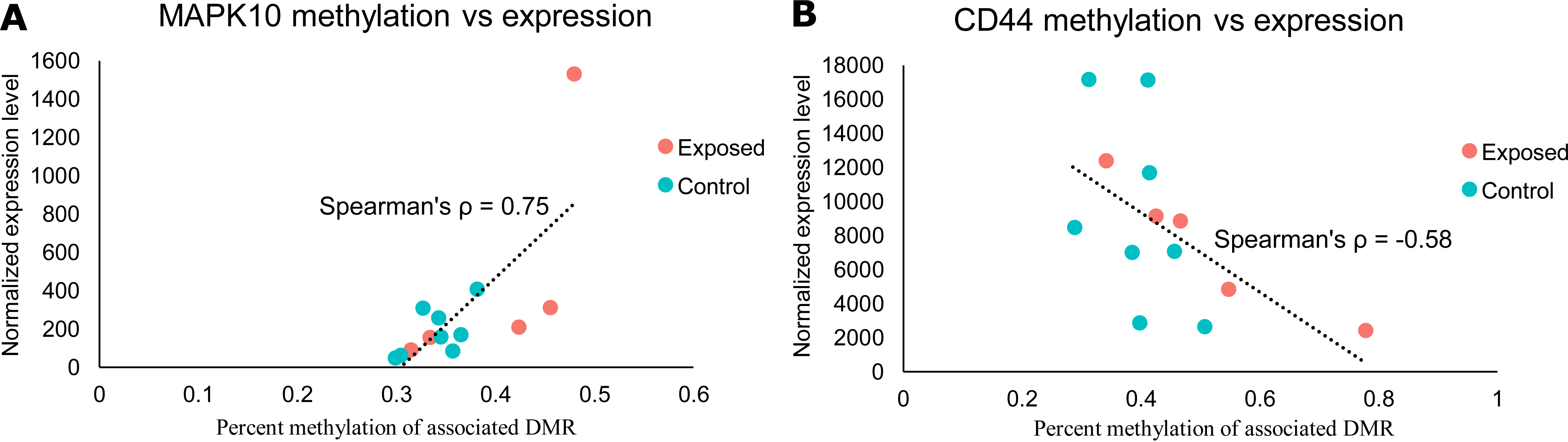
Correlation plots between expression and methylation for A) MAPK10and B) CD44. Each individual point represents one sample. Expression and methylation were significantly correlated (spearman p-value < 0.05) for both genes.

## DISCUSSION

Utilizing rhesus macaques that experienced the harsh conditions of the 2008 California wildfire season in their first three months, we have elucidated some of the long-term effects of early-life exposure to wildfire smoke. Baseline methylation profiles generally clustered better by exposure status than expression profiles (Supplementary Figure 3). Many genes (2139) were associated with differentially methylated regions between exposed and control macaques (empirical p < 0.05), while only 1 gene (*FLOT2*) was differentially expressed between these groups after multiple hypothesis correction (FDR < 0.05). Out of the genes associated with differentially methylated regions, 172 had methylation levels that significantly correlated with expression levels across samples, indicating that the overall epigenetic regulatory landscape ultimately led to few significant differences in baseline expression. However, the changes in DNA methylation were significantly enriched at promoters and enhancers, and located at regions that transcription factors may bind, suggesting that they may have an impact on gene regulation.

*FLOT2* (flotillin 2) encodes a caveolae-associated, integral membrane protein that belongs to the lipid raft family. Flotillins are implicated in variety of cellular functions, including regulation of G-protein coupled receptor signaling (20), endocytosis (21), cell-cell adhesion (22), uropod formation and migratory capacity of neutrophils and monocytes (23) and T cells (24).

*FLOT2* also protected lung epithelial cells from Fas-signaling mediated apoptosis (25), and silica nanoparticles were found in Flotillin-1 and -2 marked vesicles in alveolar epithelial cell (26). However, its role in response to wildfire smoke exposure has not been reported. One potential explanation for the few gene expression changes despite more widespread methylation differences is that many of these DMRs are in regions associated with bivalent chromatin marks. The differential methylation at these regions may not affect gene expression because the bivalent chromatin marks generally keep expression repressed, but poised for rapid activation during early development (27) or in cancer (28). This would imply that some of the methylation differences were due to early-life events that were not reflected in baseline transcript levels later in life. Additionally, although baseline gene expression was relatively similar between exposed and control macaques, one hypothesis is that the altered regulatory landscape could lead to differences in expression upon additional immune (or other) challenges. This hypothesis is supported by a previous study on macaques from these same cohorts that found differences in IL-6 (significant in males) and IL-8 (significant in females) production in peripheral blood mononuclear cells (PBMC) from wildfire smoke-exposed macaques compared to controls after a challenge with media, LPS, or flagellin (11). Out of 84 genes tested, only two (RELB and REL) showed significant differences in expression following a media challenge (essentially a comparison of baseline expression), while five genes were differentially expressed following a challenge with either LPS or flagellin (11). RELB was the only gene that was differentially expressed in all three tests, but the direction of change in challenged cells (increased RELB in cells from exposed animals) was opposite of what was found at baseline (decreased RELB in cells from exposed animals) (11). In summary, there were very few differences in baseline expression in the previous study between exposed and control cells, and even when there was differential expression, those patterns changed or became non-significant following an immune challenge. While the sample types (PBMCs vs. nasal epithelium) and ages of the macaques (adolescents vs. adults) differ between the prior study and the current study, they both support that early exposure to wildfire smoke did not lead to drastic differences in baseline expression profiles between samples. Another potential explanation for differences in the degree of differential expression and differential methylation is that we had fewer samples for our differential expression analysis, potentially limiting our ability to identify differential expression compared to our ability to identify differential methylation. If there were widespread differences in expression due to exposure status, however, we expect that wildfire smoke-exposed samples would have clustered together in the principal component analysis and in hierarchical clustering analyses, so we postulate that this is not the major reason for the lack of differential expression.

### Long-term effects of wildfire smoke exposure on the methylome

Our data implies that there are long-term effects on the methylome due to wildfire smoke exposures during infancy. DMRs were enriched for many pathways linked to asthma, COPD, or other pulmonary diseases, including *IL-15 production* (29, 30), *CXCR4 signaling* (31, 32), *Actin cytoskeleton signaling* (33, 34), *VDR/RXR activation* (35, 36), *Th1 and Th2 activation pathway* (37, 38), and *Wnt/β-catenin signaling* (39, 40) (Supplementary Table 2). Cytokines derived from T helper type 2 (Th2) cells have long been thought to play a critical role in allergic asthma through regulation of immunoglobulin E (IgE) synthesis (38, 41), but other T helper subsets (such as Th1) are starting to gain recognition for their role in asthma as well. Increased levels of the Th1 cytokine IFN-γ have been shown to exacerbate existing asthmatic responses (42) and increase airway hyperresponsiveness (41) in transgenic mice. IFNGR2 (interferon gamma receptor 2) was differentially methylated in our comparison (as were several other Th1 related genes, including IL6R, LOC694631/IFNA1/13-like, and NFATC1), perhaps indicating that the early life wildfire smoke exposure has altered Th1 responses and resulted in differential responses to bacterial and viral infection. Additionally, hypermethylation of IL6 and IFNA13 was associated with idiopathic pulmonary fibrosis (IPF) (43), while hypermethylation of IL6R was associated with COPD in prior studies (44). IL6R and IFNA13 were also hypermethylated in exposed macaques in our current study (Supplementary Table 11), indicating that changes in the Th1 pathway may contribute to the reduction in lung function noted in macaques exposed to wildfire smoke early in life (11). Th2-related genes that were differentially methylated in our dataset include IL4R and TIMD4, while there were several genes associated with DMRs that were related to both the Th1 and Th2 pathways (including CD4, IL10, IL12RB2, NFATC2, RUNX3, and SOCS3). Hypermethylation of NFATC2 (44), RUNX3 (44), and SOCS3 (45) has been associated with COPD (Supplementary Table 11). These three genes were also hypermethylated in wildfire smoke-exposed macaques versus controls.

Deletion of Fra1, the transcription factor with the most enriched motif in the DMRs (Table 2, Supplementary Table 5), in mice led to greater levels of progressive interstitial fibrosis (46). Fra1 is a bZIP transcription factor and bZIP transcription factor binding is generally inhibited by methylation (15, 16). Meanwhile, overexpression of Fra2 (another highly enriched bZIP TF motif in the DMRs) in mice lead to non-allergic asthma development (47). The other bZIP transcription factors whose motifs were among the top ten enriched motifs have also all been linked to pulmonary disease (ATF3 (48), JunB (49), BATF (50), and AP-1 (46, 51)). The role of bZIP transcription factors in pulmonary disease pathogenesis combined with the sensitivity of bZIP to changes in methylation imply that the differences in methylation noted between wildfire smoke-exposed and non-exposed macaques could greatly impact how bZIP targets are regulated following a respiratory challenge. Overall, differences in methylation in Th1 and Th2-related genes (and the relation of those genes to asthma, IPF, and COPD pathogenesis; see Supplementary Table 11) may explain the long-term differences in lung function previously observed between wildfire smoke-exposed macaques and controls (11).

Interestingly, there is also recent research that suggests that exposure to air pollution can have negative neuropsychological effects in children (52, 53). The DMRs from our dataset were enriched in multiple IPA neurological pathways, including *axonal guidance signaling* (most significant pathway), *synaptogenesis signaling pathway* (third most significant pathway), and *neuropathic pain signaling in dorsal horn neurons* (Supplementary Table 2). Additionally, the top enriched biological process term in GOfuncR (54) was *neuron differentiation*, while the top enriched cellular component term was *synapse* (Supplementary Figure 5). The effect of wildfire smoke on neurological development is understudied, but studies have shown that particles less than 0.1 µm in diameter (which are produced by wildfires) can cross the blood-brain barrier (55). Additionally, exposure to these ultrafine particles has been associated with ADHD, autism, and declines in school performance and memory in children (53). Along with this evidence from prior studies, the differential methylation of regions near genes involved in neurological pathways indicates that early-life wildfire smoke exposure could have a long-lasting impact on nervous system function.

### Genes with correlated changes in methylation and expression are enriched for pathways associated with respiratory diseases

In addition to directly studying genes and enriched pathways associated with DMRs, we also wanted to identify genes that showed correlations between expression and methylation to get a better understanding of how differences in methylation modify mRNA expression. Though only one gene was differentially expressed between our groups following multiple hypothesis correction (*FLOT2*), there were many more genes associated with DMRs that had a significant correlation between methylation and expression (172 in total; Supplementary Table 9). *MAPK10* (Spearman’s ρ ∼ 0.75) and *WNT8B* (Spearman’s ρ ∼ 0.82) were two other genes that were associated with DMRs that showed a significant correlation between methylation and expression (Supplementary Table 9). Wnt signaling has been linked to *in utero* lung development and development/maturation during early life (alveologenesis) (56–58). Prior studies have shown that Wnt/β-catenin and the mitogen-activated protein kinase (MAPK) signaling pathway take part in the airway remodeling process in asthma (39). In a mouse model of asthma, blocking Wnt signaling reduced airway remodeling, while p38 MAPK expression was increased in asthmatic mice compared to controls (39). MAPK10 expression was slightly higher on average in wildfire smoke-exposed macaques than control macaques, and methylation was significantly positively correlated with expression (hypermethylated in exposed animals). WNT8B expression was slightly lower on average in exposed macaques, while methylation was significantly negatively correlated with methylation (hypomethylated in exposed animals). Neither of these genes were significantly differently expressed, however. Given the role of Wnt signaling and MAPK signaling in airway remodeling, it seems possible that changes in gene regulation could have contributed to the reduced lung function noted in wildfire smoke-exposed macaques (11).

Our study had several limitations. As previously touched upon, our current study included only female rhesus macaques, but a prior study with these macaques noted significant sex-specific differences in PBMCs challenged with LPS or flagellin. Male wildfire smoke-exposed macaques had significantly higher levels of IL-8 compared to controls, while female wildfire smoke-exposed macaques had significantly higher levels of IL-6 compared to controls (11). While IL-6 was not differentially expressed or methylated in our exposed macaques compared to controls, this does underscore that we may have missed some sex-specific differences in gene expression or methylation by sampling only female macaques for our current study. Indeed, studies have shown that there are sex-specific differences in expression between female and male asthmatics (59, 60), implying that the molecular underpinnings of asthma and other pulmonary issues may differ between the sexes. Additionally, our cohort of wildfire smoke-exposed macaques was roughly one year older than our cohort of control macaques. Studies have indicated that methylation patterns are associated with aging (epigenetic clocks) in humans (61, 62), so this is likely the case for rhesus macaques as well. Out of 2139 genes that were associated with DMRs in our dataset, 20 were differentially methylated in a pattern that was consistent with the models from the previously referenced studies on epigenetic clocks. Based on these results, most of the differential methylation we observed cannot be explained by known differences in how methylation correlates with age. Another potential alternative explanation is that the differences in methylation we observed were due to greater cumulative exposure to pollutants in the older macaques. Table 1 shows that the difference in cumulative exposures to high levels of PM_2.5_ and ozone between the two groups were roughly equivalent to the differences observed in the first three months of life, implying that these early exposures were key drivers of the noted differences between the groups. However, cumulative exposures below the current U.S. EPA standards were associated with increased mortality in a Medicare population (63), and they may also have an impact on the epigenome. The epigenetic effects of acute and chronic wildfire smoke exposure are worthy of further investigation. As previously discussed, we had a smaller sample size for our expression dataset (n = 13 after removing two outliers) than our methylation dataset (n = 22). This could explain why we saw fewer changes in expression overall, however samples appeared to cluster more closely based on exposure status for the methylation dataset than the expression dataset (Supplementary Figure 3). The p-values from DMRichR were empirical p-values calculated from permutation tests (64, 65). Although this puts our study at a higher risk of false-positive findings, these permutation p-values calculated by DMRichR were used to determine DMR significance in multiple published studies in combination with effect size (64–67). Given that our analysis of chromatin states relied on human hg38 annotations, we compared our macaque rheMac10 annotations to the hg38 annotations to make sure they were similar enough. About 67% of the DMR gene annotations were exact matches after lifting over the coordinates to hg38. While 30% of DMRs had a different annotation, some of these differences were just due to differences in gene naming convention between the species. For example, one DMR was annotated to LOC694631 (IFNA1/13-like) in rheMac10, while the lifted over DMR was annotated to IFNA13 in hg38. In a broader pathway analysis, 90% of the IPA pathways enriched in DMRs using rheMac10 annotations were also enriched when we used hg38 annotations. We also focused our discussion on genes that were consistent between the two annotations.

One area of interest for future studies would be the stability of these changes. The exposure event took place in 2008, while samples were collected from the macaques in 2019. Over that relatively long course of time (the average lifespan for macaques in captivity is ∼27 years (68)), the methylation profiles still clustered based on exposure status (Figure 2, Supplementary Figure 3). This implies that there are long-term impacts of wildfire smoke exposure on methylation, and that at least some of these changes are highly stable. An early study on DNA methylation stability involved sampling individuals three days apart to check for differences in DNA methylation. This study on 12 gene promoters indicated that methylation stability was marker dependent and varied based on sequence composition (69). Meanwhile, a large-scale study on how storage conditions affect methylation stability indicated that storing DNA samples in temperatures as high as four degrees Celsius for up to 20 years had no significant impact on methylation (70).

## CONCLUSIONS

In summary, our study revealed differences in methylation and gene expression in nasal epithelial samples between macaques that were exposed to wildfire smoke during early life and macaques that were not exposed to wildfire smoke during early life. The wildfire smoke associated DMRs were enriched for a variety of immune processes, but there were few significant expression differences at baseline between exposed and non-exposed macaques. Given the differences in methylation, perhaps differences in expression between these two groups would become apparent following an immune/respiratory challenge, but future studies would be required to explore this hypothesis. Our study indicates that wildfire smoke exposure in early life can have long-term impacts on the epigenome.

## METHODS

### Animals

Wildfire smoke-exposed rhesus macaque monkeys born between April 1 and June 8, 2008 were housed in outdoor facilities at the CNPRC from birth to now (Table 1). Monkeys born between April 1 and June 8, 2009 were used as controls. PM_2.5_ and ozone were measured by a California Air Resources Board air monitoring station (site no. 57,577) located 2.7 miles southeast of the California National Primate Research Center on the University of California Davis campus (Figure 1). Care and housing of animals complied with the provisions of the Institute of Laboratory Animal Resources and conformed to practices established by the American Association for Accreditation of Laboratory Animal Care. Procedures in this study were approved by the UC Davis Institutional Animal Care and Use Committee.

### Sample Collection and DNA/RNA Extraction

Nasal epithelium samples were collected from 22 female rhesus macaques (*Macaca mulatta*) housed at the California National Primate Research Center. Exposures of these animals to wildfire smoke were previously estimated (11). Eight of these macaques were born in 2008 and exposed to wildfire smoke from birth to 3 months old, while the other 14 were born in 2009 with low wildfire exposure from birth to 3 months old (Table 1, demographic comparison of two groups). We collected these nasal epithelium samples in 2019. RNA and DNA were isolated using the Allprep DNA and RNA kit (Qiagen) according to the manufacturer’s instructions.

### Library preparation for whole genome bisulfite sequencing (WGBS)

Whole genome bisulfite sequencing (WGBS) libraries were prepared for all 22 samples. Library quality was checked prior to sequencing using an Agilent 2100 Bioanalyzer system; library concentration was measured using a Qubit DNA high sensitivity assay. Each library was comprised of sample from a single individual; these individually barcoded libraries were then pooled and sequenced on two lanes from a NovaSeq 6000 S4 flow cell at PE150 using Swift’s Accel-NGS Methyl-Seq Kit at the DNA Technologies and Expression Analysis Cores at the UC Davis Genome Center. We sequenced approximately 475 million paired end reads per sample that passed initial filters. Reads were demultiplexed using the bcl2fastq Illumina software.

### WGBS read alignment, differential methylation analysis, pathway analysis, and chromatin state analysis

The CpG_Me pipeline (71–74) was utilized to align the WGBS data. Reads were trimmed using Trim Galore (73) to address methylation biases at the 5’ and 3’ end of reads (10 bases were trimmed from the 3’ end of both read 1 and read 2, and 10 and 20 bases were trimmed from the 5’ end of reads 1 and 2 respectively). The reads were aligned to the *M. mulatta* genome using Bismark (72), which was also used to deduplicate the aligned reads and generate CpG count matrices. Read quality and mapping quality were assessed using MultiQC (74). Differentially methylated regions between exposed and non-exposed macaques were identified using DMRichR (64, 75, 76), which uses the dmrseq (75) and bsseq (76) algorithms. Animal weight was adjusted for as a covariate. We used the default paramters for DMRichR, including requiring at least 1x coverage for all samples for a CpG, requiring a minimum of 5 CpGs for a DMR, performing 10 permutations for DMR and block analyses, and setting the single CpG coefficient required to discover testable background regions to be at least 0.05. Using DMRichR, candidate regions are identified based on differences in mean methylation between groups, then region-level metrics that account for mean methylation, CpG correlation, and coverage are computed. These region-level metrics are then compared to a pooled null distribution generated via permutations to calculate an empirical p value for each candidate region (64, 65). Bsseq (76) was used to generate individual smoothed methylation values and heatmap visualizations. IPA (QIAGEN Inc., https://www.qiagenbioinformatics.com/products/ingenuitypathway-analysis) was used for pathway enrichment analysis. We also used GOfuncR (54) for GO enrichments based on DMR coordinates rather than gene names. HOMER (13) was used to identify enriched transcription factor binding motifs in the DMRs (p < 0.05), while we utilized MethMotif (14) to characterize methylation frequency of transcription factors whose binding motifs were enriched in the DMRs. We used the UCSC liftover tool (77) to lift DMR coordinates from rheMac10 to hg38 because chromatin state information was not available for *M. mulatta*. Locus Overlap Analysis (LOLA) (78) was used to determine whether DMRs were enriched for chromHMM (79) states relative to the background regions. The spearman correlation coefficient between gene expression and methylation levels for genes associated with DMRs was used to determine whether significant methylation changes were associated with changes in gene expression (p< 0.05).

### Library preparation for RNA-seq

RNAseq libraries were prepared for a total of 15 samples: 6 from wildfire smoke-exposed individuals and 9 from non-exposed individuals (Supplementary Table 12, comparison of these two groups). As some of the RNA samples were of low quantity, a special low-input RNA-seq pipeline were applied at the Genomics, Epigenomics and Sequencing Core of University of Cincinnati (80, 81). Briefly, polyA RNA was isolated using NEBNext Poly(A) mRNA Magnetic Isolation Module (New England BioLabs, Ipswich, MA) and enriched using SMARTer Apollo NGS library prep system (Takara Bio USA, Mountain View, CA). Libraries were prepared using NEBNext Ultra II Directional RNA Library Prep Kit (New England BioLabs), indexed, pooled and sequenced using Nextseq 550 sequencer (Illumina, San Diego, CA). Approximately 40 million reads passing filter per sample were generated under the sequencing setting of single read 1x85 bp. Reads were demultiplexed and adapters were trimmed using the bcl2fastq Illumina software.

### RNA-seq read alignment, differential expression analysis, pathway analysis, and co-expression analysis

Read quality was checked using *FastQC* (82), then the reads were aligned to the *Macaca mulatta* genome (rheMac10, GenBank assembly accession: GCA_003339765.3) with *Bowtie2* (83). Transcripts were quantified using *RSEM* (84). The data from *RSEM* was congregated and converted into *DESeq2* (85) format using *tximport* (86). Sample clustering by expression (investigated via principal component analysis and hierarchical clustering) and detection of differentially expressed genes between wildfire smoke-exposed and non-exposed samples was done using *DESeq2* (85). Individual weight was included as a covariate in the differential expression analysis. Two samples (one wildfire smoke-exposed and one non-exposed sample) were excluded from all subsequent RNA-sequencing analyses because they were identified as outliers in the hierarchical clustering analysis (Supplementary Figure 6). The resulting log-fold change values were shrunken (following the recommendation from the *DESeq2* reference manual) using *apeglm* (87). Differentially expressed genes had FDR < 0.05 and an absolute shrunken fold change of at least 1.2. The Ingenuity Pathway Analysis (IPA) software (QIAGEN Inc., https://www.qiagenbioinformatics.com/products/ingenuitypathway-analysis) was used for pathway analysis. Significantly enriched pathways in IPA had a p-value < 0.05.

Co-expressed modules of genes were found using WGCNA (19). The soft threshold (power) was set to 8 based on a plot of soft threshold vs scale free topology model fit. Modules that were too similar to one another (below a height of 0.5) were merged into one module. After merging, the final co-expression modules were tested for significant associations with wildfire smoke exposure and animal weight. Pathways enriched in genes in modules of interest were identified using IPA.

## DECLARATIONS

### Ethics approval and consent to participate

Procedures in this study were approved by the UC Davis Institutional Animal Care and Use Committee.

### Consent for publication

Not applicable.

### Availability of data and materials

WGBS and RNA-seq data will be deposited to GEO upon manuscript acceptance.

### Competing interests

The authors declare they have no competing interests.

### Funding

This study was supported by a pilot grant from Environmental Health Sciences Center (NIEHS-P30 ES006096) awarded to HJ. HJ was also supported by an Environmental Health Sciences Scholar Award (P30 ES006096), and NIH/NIAID R01AI141569-1A1. BL was supported Canadian Institutes of Health Research (CIHR) postdoctoral fellowship (MFE-146824) and a CIHR Banting postdoctoral fellowship (BPF-162684).

### Authors’ contributions

HJ conceived the study in discussion with LAM and JML. APB drafted the manuscript with the help of HJ, BIL, JML and LAM. LC extracted DNA and RNA from nasal samples, prepared WGBS libraries. APB performed WGBS analysis with the help of BIL. APB performed other data analysis and visualization in discussion with HJ. All authors have approved the final version of this manuscript.

## Acknowledgements

Nasal sampling was performed by staff members at CNPRC Research Services. The sequencing was carried out at the DNA Technologies and Expression Analysis Cores at the UC Davis Genome Center, supported by NIH Shared Instrumentation Grant 1S10OD010786-01.

---

**Supplementary Table 1.** Differentially methylated regions between rhesus macaques exposed to wildfire smoke in early life and rhesus macaques with no early life exposure to wildfire smoke.

Note: betaCoefficient is presented with respect to exposed macaques (*i.e.* a positive betaCoefficient implies hypermethylation in exposed macaques compared to control macaques)

**Supplementary Table 2**. Canonical pathways from Ingenuity Pathway Analysis that were enriched in all differentially methylated regions.

**Supplementary Table 3.** Canonical pathways from Ingenuity Pathway Analysis that were enriched in differentially methylated regions hypermethylated in macaques exposed to wildfire smoke in early life.

**Supplementary Table 4.** Canonical pathways from Ingenuity Pathway Analysis that were enriched in differentially methylated regions hypomethylated in macaques exposed to wildfire smoke in early life.

**Supplementary Table 5.** Transcription factor binding site motifs that were significantly enriched in differentially methylated regions (from HOMER (1)).

**Supplementary Table 6.** The differentially expressed gene between rhesus macaques exposed to wildfire smoke in early life and rhesus macaques with no early life exposure to wildfire smoke.

Note: log2FoldChange is presented with respect to exposed macaques (*i.e.* a positive log2FoldChange implies greater expression in exposed macaques compared to control macaques).

**Supplementary Table 7.** Genes in the purple module (the module most significantly associated with exposure) from the weighted gene coexpression network analysis (WGCNA (2)).

**Supplementary Table 8.** Canonical pathways from Ingenuity Pathway Analysis that were enriched in genes in the purple module from the WGCNA (2) analysis.

**Supplementary Table 9.** Genes that showed significant correlation (p ≤ 0.05) between methylation and expression across all samples.

**Supplementary Table 10.** Canonical pathways from Ingenuity Pathway Analysis that were enriched in genes that had significantly correlated methylation and expression.

**Supplementary Table 11.** Comparison between differentially methylated genes from the current study and other studies on respiratory diseases.

**Supplementary Table 12.** Extended information on the samples in our current study.

**Supplementary Figure 1.**
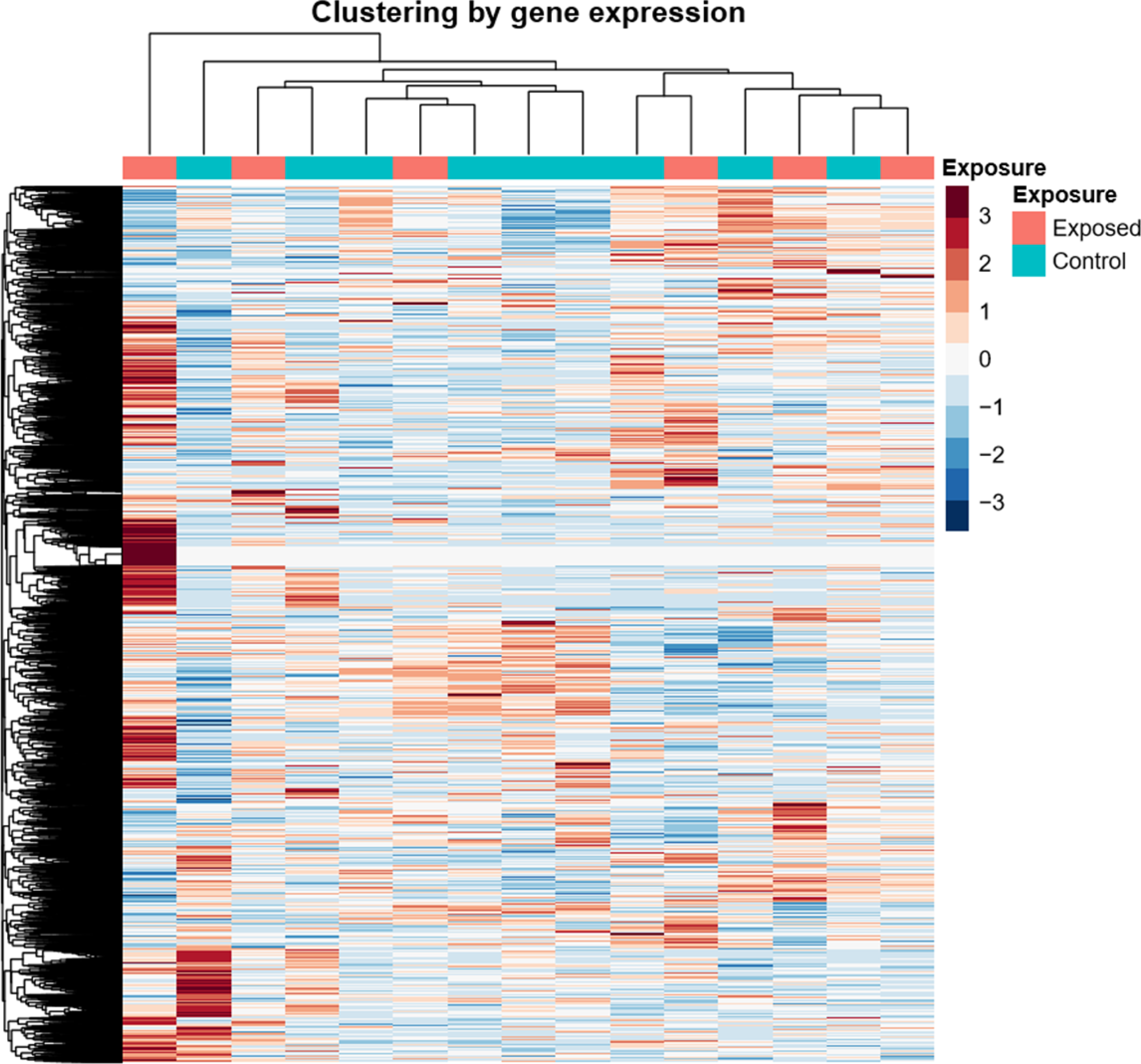
Enrichment of different CpG features associated with all differentially methylated regions, regions hypermethylated in wildfire-exposed macaques, and regions hypomethylated in wildfire-exposed macaques. Asterisks indicate a significant deficit or enrichment of the feature in a given set (p ≤ 0.05).

**Supplementary Figure 2.**
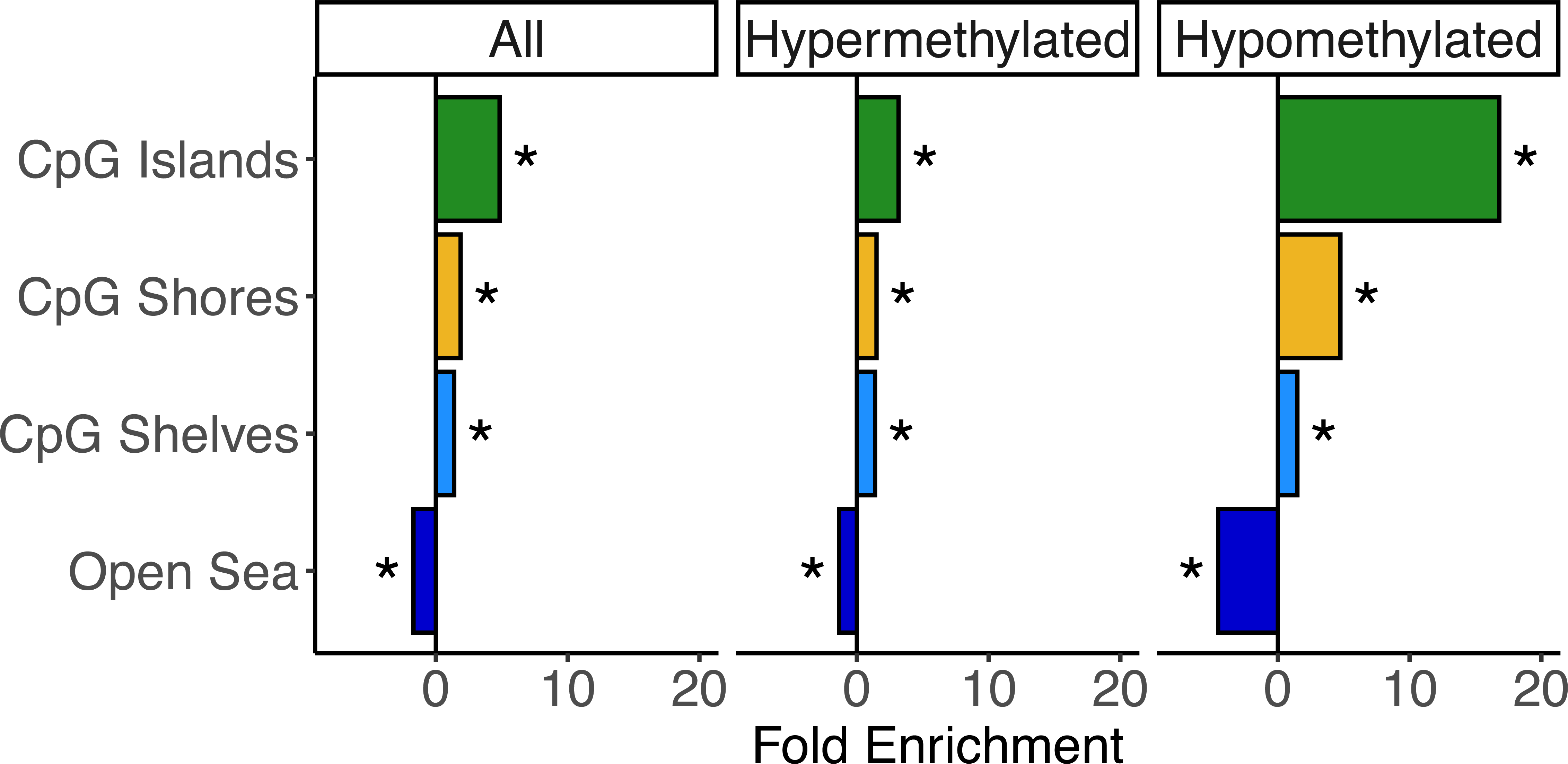
Enrichment of different genic features associated with all differentially methylated regions, regions hypermethylated in wildfire-exposed macaques, and regions hypomethylated in wildfire-exposed macaques. Asterisks indicate a significant deficit or enrichment of the feature in a given set (p ≤ 0.05).

**Supplementary Figure 3.**
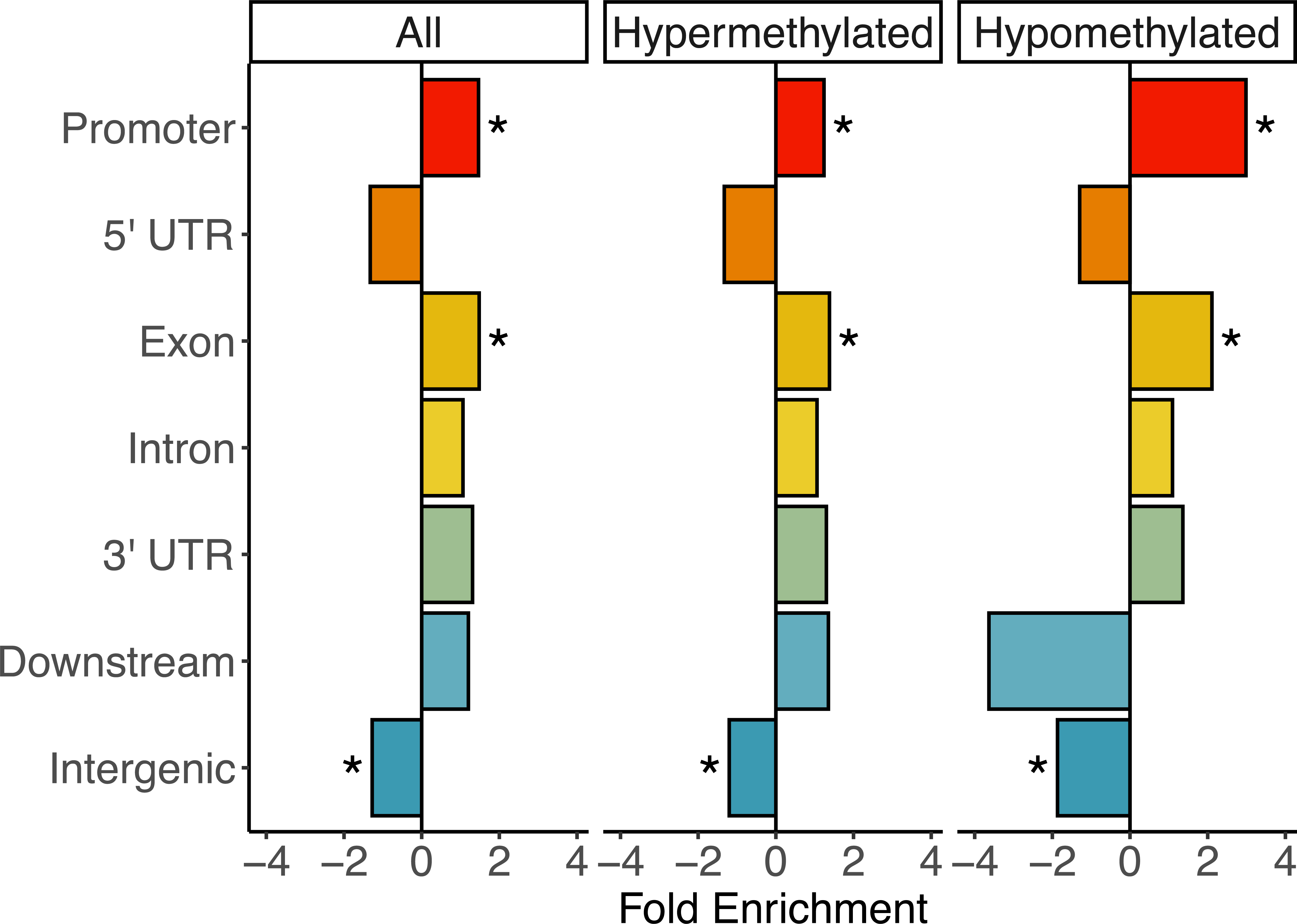
Heatmaps showing sample clustering by A) methylation and B) gene expression, and principal component analysis showing sample clustering by C) methylation and D) gene expression.

**Supplementary Figure 4.**
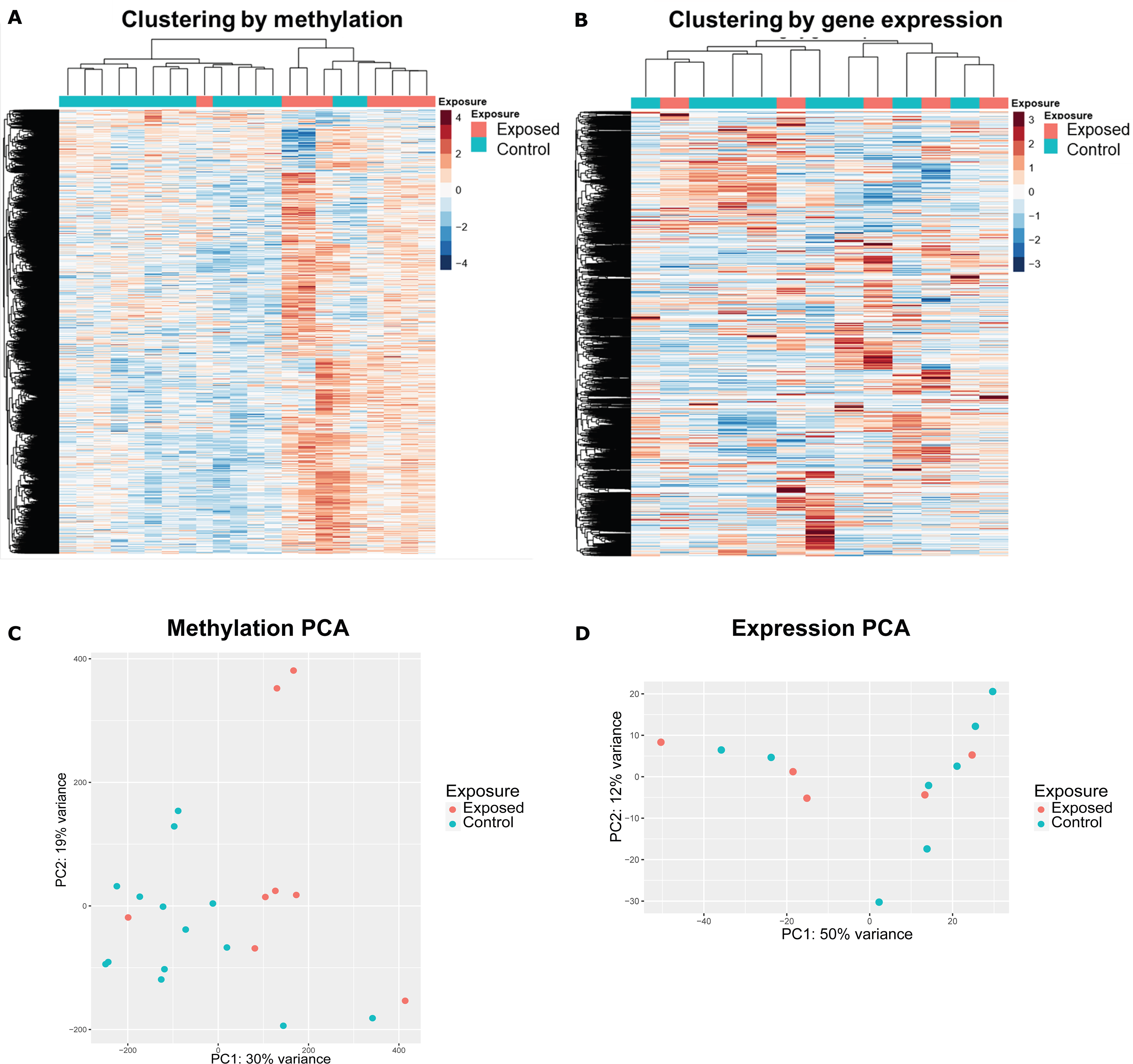
Module-trait relationship between clusters identified in WGCNA and either exposure or animal weight. The top number in each box is the correlation value (ranging from -1 to 1), while the bottom number in parentheses is the p-value for this correlation.

**Supplementary Figure 5.**
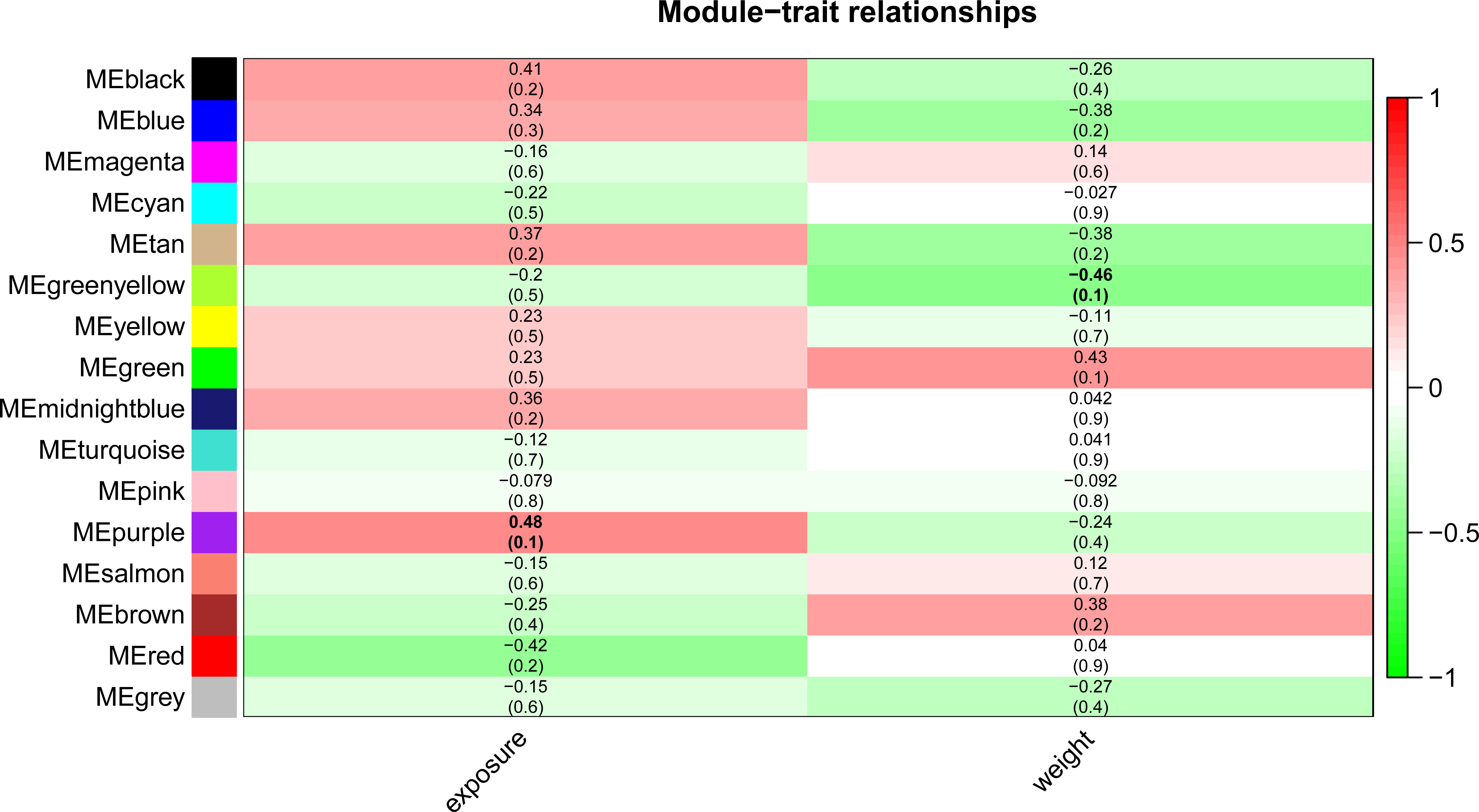
Top enriched biological process, cellular component, and molecular function gene ontology terms identified by GOfuncR (3) associated with differentially methylated regions between wildfire-exposed macaques and control macaques.

**Supplementary Figure 6.**
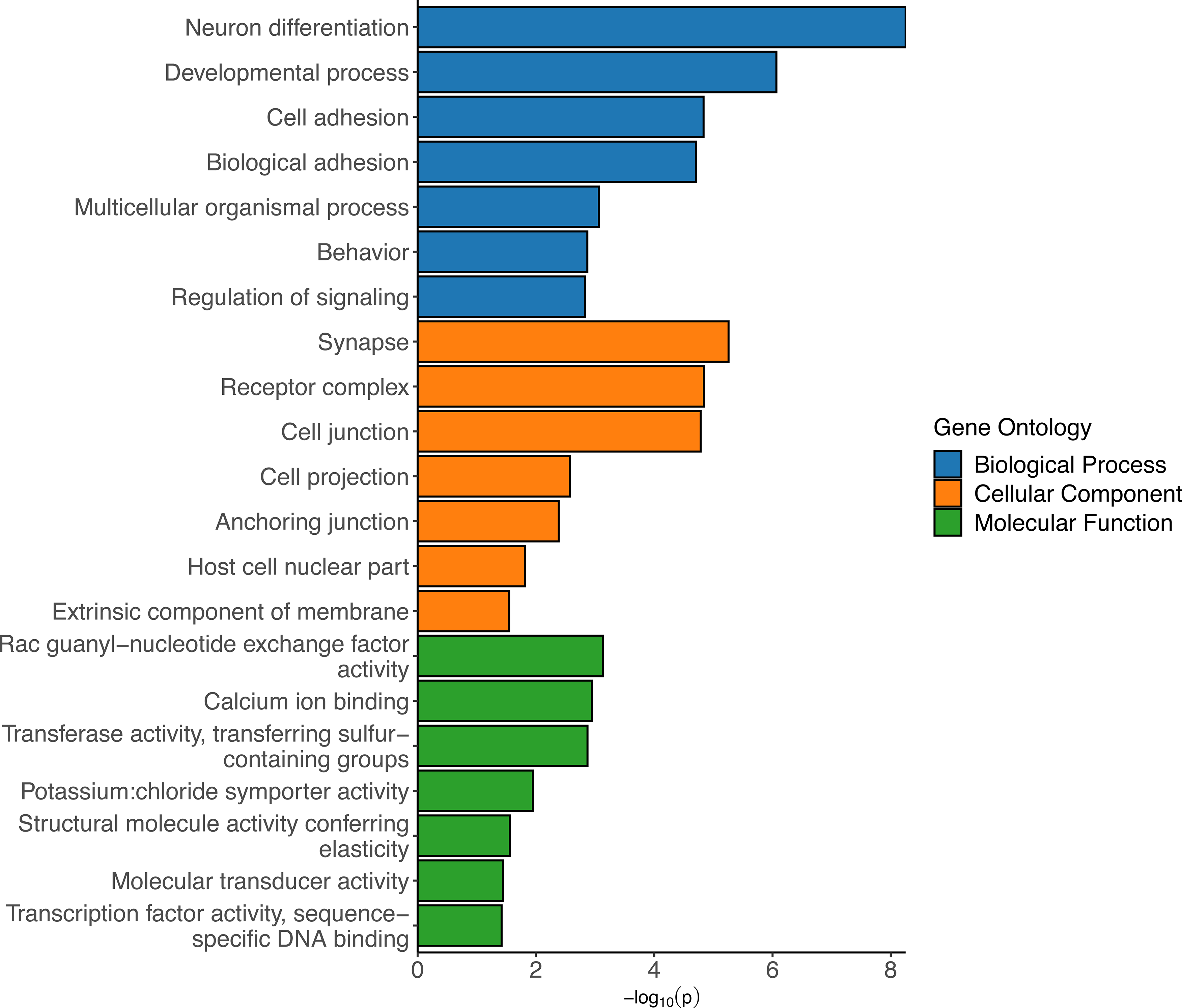
Heatmap showing all samples clustering by gene expression. Gene expression data from the two leftmost samples were removed from the study as outliers.

## Notes

### Competing Interest Statement

The authors have declared no competing interest.

